# A chromosome-level genome assembly of *Dracaena cambodiana* and comparative genomics analysis highlights the distinct biological features of two resource species of dragon’s blood

**DOI:** 10.1101/2024.07.03.601834

**Authors:** Yanhong Xu, Junxiang Peng, Sipeng Li, Yang Liu, Dong Wen, Jiemei Jiang, Mei Rong, Wei Wei, Qiuling Wang, Yue Jin, Shuwen Yang, Siyu Wang, Jian-he Wei

## Abstract

This study reports a chromosome-level genome for *Dracaena cambodiana*, one of two typical dracaena species in China. This work will help to deepen the understanding of the dracaena species and the conservation and development of plant resources. The final assembly consisted of 54 scaffolds, spanning 1.08 Gb, with a scaffold N50 size of 52.29 Mb, encoded 36860 predicted protein-coding genes. A total of 1,064,810,157 bp of genome sequences were located on 20 chromosomes (2n = 40), accounting for 98.46%. We investigate the possible mechanisms of the longevity of dracaena, the longest-lived plant in the world, which involve DNA repair and post-translational modification. We also performed comparative genomic analysis of the previously assembled genome of *Dracaena cochinchinesis* with this genome, and found that the two involved interaction of plant−pathogen interaction and ubiquitin mediated proteolysis, which may reveal why Dracaena has environmental adaptability and longevity characteristics.

## Introduction

Dragon’s blood, a precious traditional medicine, has been widely used in the world since ancient times. It has the effects of promoting blood circulation and removing blood stasis, astringent hemostasis, anti-inflammatory and analgesic. It has many pharmacological activities, such as promoting blood circulation, hemostasis, anti-inflammatory, analgesic, anti-dysentery, antidiarrheal, anti-ulcer, antimicrobial, anti-tumor and so on (Gupta et al., 2008; Luo et al., 2011; Fan et al., 2014; Sun et al., 2019; Maděra et al., 2020; Liu et al., 2021; Liu et al., 2022;). The original plants of dragon’s blood are some species of *Dracaena* spp., such as *Dracaena cinnabari* from sokotra island in Yemen and *Dracaena draco* from West Africa and India, which regarded as the earliest origin members of the genus *Dracaena* (Gupta et al., 2008; Liu et al., 2021). With the discovery of substitutes, the term “Dragon’s blood” in general is used for all kinds of resins and saps obtained from four distinct plant genera, including *Croton* (Euphorbiaceae), *Dracaena* (Dracaenaceae), *Daemonorops* (Palmaceae), and *Pterocarpus* (Fabaceae) (Gupta et al., 2008). The genus *Dracaena* Vand. ex L. is placed in the family Asparagaceae subfamily Nolinoideae (Lu et al., 2014; Maděra et al., 2020). There are more than 60–190 species of the genus *Dracaena* in the world (Bos,1998; Mabberley, 2008; Govaerts et al., 2019) and mainly distribution in tropical and subtropical regions of Asia and Africa. According to Flora of China, there are 5 *Dracaena* species in China, of which *D. cochinchinensis* and *D. cambodiana* were the mainly plant resources of dragon’s blood.

It is believed that *Dracaena* is long-lived and slowly growing species, and only trees that are over 30 years of age have been shown to produce dragon’s blood (Edwards et al., 1997; Tomlinson and Huggett, 2012; Ding et al., 2020; Maděra et al., 2020; Xu et al., 2022). Additionally, only in response to external stimuli, dragon’s blood is secreted, while injury in nature is a random event. The contradiction between limited availability and market demand caused overexploitation of the wild resources leading to severe depletion of the species, and some species are listed in the IUCN Red List (IUCN red list, 2017). In China, *D. cochinchinensis* and *D. cambodiana* are listed as the second-class state-protected plants in the National Key Protected Wild Plants of China and prohibited logging (Hubálkováet al., 2015; Kamel et al., 2018; Xu et al., 2022).

Dragon trees are famous in the world, not only for their secretion of red resin, but most of them have high values in gardening and horticulture as a green plant. In China, *D. cochinchinensis* is the official original species found to produce dragon’s blood and the Pharmacopoeia of the People’s Republic of China recorded *D. cochinchinensis* (Lour.) S.C.Chen as the only source plant (Zheng et al., 2009; Luo et al., 2011; Sun et al., 2021; Xu et al., 2022). *D. cambodiana* is also included in the production standards in some region and as another species used in China as source of dragon’s blood (Yang et al., 2018). Moreover, *D. cambodiana* has high ornamental value in horticultural greening due to its beautiful tree shape. One problem is that the name of two species in *Dracaena* genus is still debated. According to the latest report, *Aletris cochinchinensis* Lour. and *Dracaena cochinchinensis* (Lour.) S. C Chen is the same species and should be reduced as a synonym for *Dracaena angustifolia* Roxb. It is also found *D. cambodian* in China were divided into two independent branches, namely the Yunnan branch (*D. cambodian* A) and the Hainan branch (*D. cambodian* B), and the Hainan branch (*D. cambodian* B) is suggested to be a hidden specie to be published or a new record specie in China (Xin et al., 2024).

The main components of dragon’s blood are varieties of flavonoids, most of which are based on 7,4’-dihydroxyflavan as the basic skeleton (Sun et al., 2021). Oligomeric flavonoids, composed of one dihydrochalcone unit condensed with one or more chalcone, flavane, or homoisoflavane units, account for over 50% of the resin produced by *Dracaena* spp.(Hao et al., 2015; Pang et al., 2016). Thus far, hundreds of flavonoids have been isolated from the resin wood of *D. cochinchinensis* (Gupta et al., 2008; Fan et al., 2014; Sun et al., 2019; Sun et al., 2021; Xu et al., 2022). Especially 4’-hydroxy-2,4-dimethoxydihydrochalcone (loureirin A) and 4’-hydroxy-2,4,6-trimethoxydihydrochalcone (loureirin B) have been used as chemical markers for quality control of dragon’s blood (Sun et al., 2021). In addition, terpenoids and steroidal saponins are also effective components with antitumor and cytotoxic activity (Maděra et al., 2008). Though there have been many studies on the chemical composition and pharmacological effects of dragon’s blood (Gupta et al., 2008; Fan et al., 2014; Sun et al., 2019), the genetic mechanism of dragon’s blood formation has not been explored to a large extent. Research has shown that a large amount of flavonoids begin to synthesize after 3 days of wounding stress induction, and the expression of key enzyme genes in the flavonoid biosynthesis pathway is significantly upregulated (Zhu et al., 2016; Sun et al., 2021; Xu et al., 2022). With the first completion of genome sequencing of the *D. cochinchinensis*, some biological characteristics of *Dracaena* have been revealed. Meanwhile, enzyme genes and regulatory genes related to the flavonoid synthesis were systematically identified, and those may be involved in dragon’s blood synthesis pathway were predicted, which preliminarily deciphered the molecular mechanism of wound induced dragon’s blood formation (Xu et al., 2022). However, there are still big gaps in the botanical and genetic research on *Dracaena*, such as the current limited understanding of the basic plant physiology, the lack of genetic and environmental mechanisms underlying the production and chemical composition of *Dracaena* species resin, as well as the process of the red resin secretion, and so on.

So far, only the reference genome of *D. cochinchinensis* in *Dracaena* genus has been published (Xu et al., 2022). Here, we report the reference genome of *D. cambodiana* (Hainan) at the chromosome level, and conducted comparative genomics analysis with that of *D. cochinchinensis*. Our findings represent an important resource that provide further insight into the fundamental biology and the molecular basis of *Dracaena* genus, allowing an in-depth study of dragon’s blood formation, ensuring its sustainability, and contributing to the conservation of this valuable species.

## Result

### Genome Sequencing, Assembly

The genome survey using a k-mer analysis (k = 19) revealed that the genome size of *D. cambodiana* was approximately 938.93 Mb, with the heterozygosity of 1.06% and the repeat sequence ratio of 53.55% (Supplemental Figure S1; Supplemental Table S1). To construct the genome sequence of *D. cambodiana*, the datasets generated by short-read Illumina sequencing, long-range PacBio sequencing, and chromatin conformation capture (Hi-C) sequencing were integrated. We obtained 88.68 Gb of short reads sequences, 81.79 Gb of long reads, and 196.32 Gb of Hi-C data (Supplemental Table S2, S3, S4). Evaluation of the genome completeness indicated 93.9% coverage of Complete BUSCOs (C) by the Benchmarking Universal Single-Copy Orthologs (BUSCO) analysis (Supplemental Table S5) (Simao et al., 2015). Additionally, 99.78% of the of Illumina clean reads that had been compared to the reference genome accounted for the total Illumina Clean Reads (Supplemental Table S6). The chromosomal interaction signal was strong (Supplemental Figure S2), indicating that the quality of Hi-C assembly was high. The final assembly consisted of 54 scaffolds, spanning 1.08 Gb, with a scaffold N50 size of 52.29 Mb (Supplemental Table S7). A total of 1,064,810,157 bp of genome sequences were located on 20 chromosomes (2n = 40), accounting for 98.46% (Supplemental Table S8, Figure S1).

### Genome Annotation

Based on de novo and homology-based predictions and transcriptome data, a total of 36860 protein-coding genes were predicted, with an average cds length of 1,003.93 bp and an exon length of 348.49 bp (Supplemental Table S9). A total of 33506 protein-coding genes (90.90%) were successfully annotated by seven databases of KEGG (Kyoto encyclopedia of genes and genomes) (Kanehisa et al., 2023), GO (Gene ontology) (Ashburner et al.,2001), NR (Non-redundant protein) (Pruitt et al., 2005), Pfam (Pfam protein families) (Finn et al., 2014), and KOG (EuKaryotic orthologous groups) (Tatusov et al., 2003) (Supplemental Table S10). In addition, by comparing with the known noncoding RNA libraries, we obtained noncoding RNA information relating to the *D. cambodiana* genome, including 1450 transfer RNAs, 2533 ribosomal RNAs, 191 microRNAs, and 906 small nuclear RNAs (Supplemental Table S11).

### LTR insertion and genome size expansion

Further analysis showed that 71.86% of the *D.cambodiana* genome consists of repetitive sequences. As in many other sequenced plant genomes, long terminal repeat retrotransposons (LTR-RTs) dominated, accounting for 49.76 % of the assembly; 32.08 % of the genome sequence comprised Gypsy elements (346,930,526 bp), and 4.8 % comprised Copia elements (51,955,114 bp) (Supplemental Table S12).

### Genomic evolution

*D. cambodiana* belongs to the Asparagales clade of monocotyledons and is close to *Dracaena cochinchinensis*, *Asparagus officinalis*, *Dendrobium chrysotoxum*, and *Apostasia shenzhenica*, which have completed genome sequences. We clustered the annotated genes into gene families among these species along with additional monocot and dicot species, including *D. cochinchinensis, A. officinalis, D. chrysotoxum, A. shenzhenica, Elaeis guineensis, Oryza sativa, Asparagus kiusianus, Dioscorea zingiberensis, Dendrobium nobile, Dendrobium catenatum, Allium fistulosum, Vanilla planifolia* and *Amborella trichopoda*. Family clustering yielded 74129 gene families (groups), including 367 single-copy families. Evolutionary trees were constructed based on coding sequences using RAxML software. *D. cochinchinensis* and *D. cambodiana* were gathered in one branch, as were *A. officinalis* and *A. kiusianus*, thus indicating that Asparagus and Dracaena underwent independent divergence long ago (70.1 Mya); the time of divergence between *D. cochinchinensis* and *D. cambodiana* was 9.5 Mya (Figure S2).

To investigate the genome expansion in *D. cambodiana*, we analyzed whole-genome duplication (WGD) events. Ks values were estimated based on paralogous gene pairs in collinear regions detected in *D. cambodiana* and three other representative plant species (*D. cochinchinensis*, *A. officinalis*, and *A. shenzhenica*). The distribution of Ks distances in *D. cambodiana* showed two peaks at approximately 0.5 and 1.5 (Figure S3). The first peak shared by *D. cochinchinensis* and *D. cambodiana* may help both to adapt to dramatic changes in the natural environment through WGD. The second peak at approximately 0.5 in *D. cambodiana* and 1.0 in *D. cochinchinensis* indicated that these two typical dracaena tree species experienced different WGD events after experiencing a common WGD.

### Comparative genomics demonstrated the adaptation of *D.cambodiana*

Copy numbers of homologous gene families vary greatly among different species, caused by differences in rates of gene gain and loss. Analysis using Computational Analysis of Gene Family Evolution (CAFE) identified 849 expanded gene families in *D.cambodiana* compared with the common ancestor of *D.cochinchinensis* and *D.cambodiana* (Bie et al., 2006) (Figure S2). Total genes in the expanded families were annotated with GO and KEGG pathways, respectively(Supplemental Figure S3). GO analysis revealed that these expanded orthogroups were related to several metabolic processes, such as nucleic acid binding, translation, heme binding and structural constituent of ribosome. The enriched KEGG categories were ribosome, plant-pathogen interaction, phenylpropanoid biosynthesis, oxidative phosphorylation. We hypothesize that these expanded genes are closely related in some way to the functional requirements of environmental adaptability in *D.cambodiana.*

33 genes were under positive selection and showed significant enrichment in KEGG terms related to biosynthesis of cofactors, nucleotide excision repair, nucleocytoplasmic transport. Genes involved in DNA damage repair were positively selected, which may have improved the species’ ability to adapt to the environment over time.

Next, we identified unique and shared gene families among the four Asparagales species. *D.cambodiana* shared 9186 families with *D. cochinchinensis*, *D. chrysotoxum*, and *A. shenzhenica*, whereas 6455 gene families appeared to be unique to *D.cambodiana*(Figure S4). Enrichment of unique gene families through KEGG was mainly involved biosynthesis of amino acids, biosynthesis of cofactors and carbon metabolism. Through KEGG annotation analysis of the common genes of *D.cambodiana* and *D. cochinchinensis* was mainly involved RNA degradation, Plant−pathogen interaction, Ubiquitin mediated proteolysis, Protein processing in endoplasmic reticulum(Supplemental Figure S4).

## Discussion

As the original plant of the traditional medicine dragon’s blood, the germplasm resources of Dracaena are worth exploring (Liu et al., 2021). *D.cambodiana* is one of the typical Dracaena species in China, where we have performed a chromosome-level genome assembly for it, which will help us to understand the biology of this species as well as the subsequent conservation and development of resources.

The longevity of Dracaena species is of noteworthy concern. Our WGD analysis revealed that *D.cambodiana* and *D. cochinchinensis* underwent a common WGD event, which may confer a unique longevity capability to this species. And the subsequent differential WGD events may have led to the differentiated physiological traits of the two species. In addition, both have experienced positive selection on genes associated with DNA repair pathways, which would help maintain genomic stability as well as environmental adaptability (Spampinato 2017).

When injured or infested with pathogenic bacteria, Dracaena produces a red resin for defense and resistance to subsequent damage. Flavonoids are main components of dragon’s blood, and in a previous study of *D. cochinchinensis*, we identified the genes of the flavonoid synthesis pathway (Xu et al., 2022). This study in which we analyzed genes shared by *D.cambodiana* and *D. cochinchinensis*, genes related to plant-pathogen interactions were significantly enriched, which may indicate that dragon’s blood is produced in response to invasion by pathogenic bacteria. Meanwhile genes for RNA degradation as well as plant signaling pathways, MAPK, and other typical pathways in response to external stress were significantly enriched, implying that Dracaena species responds well to external stress. Ubiquitination degradation as well as endoplasmic reticulum protein processes were significantly enriched, and the degradation of error proteins generated by stress processes in Dracaena was mainly through ubiquitination. We conclude that there is a potential association between dragon’s blood production, flavonoid biosynthesis, plant-pathogen interactions, signaling pathways, and ubiquitination degradation in Dracaena species. This system contributes to a better environmental adaptation of Dracaena.

Summary, the present study reports the genome of *D.cambodiana* at the chromosome level, which provides a valuable research resource. We compared the genomes of two typical types of Dracaena at the genome level, revealing potential pathways for environmental adaptation in Dracaena. These findings help us to understand how Dracaena is adapted to the environment and the subsequent conservation development and utilization.

## Method

### Plant material, DNA extraction, and sequencing

The fresh samples from which DNA was extracted for the genome sequencing of 100-year-old *D. cambodiana* growing wildly on Boundary Island in Hainan province, China (Lingshui Li Autonomous County; N:18.576845°, E:110.201546°). The plants were formally identified by Professor Jianhe Wei.

A modified cetyltrimethyl ammonium bromide (CTAB) method was used for DNA extraction. High-quality and purified genomic DNA samples were obtained, then, DNA samples were randomly interrupted with the Covaris ultrasonic crusher. Using the Nextera DNA Flex Library Prep Kit (Illumina, San Diego, CA, USA), Illumina sequencing library with insertion fragment size of 150 bp was constructed. The genomic DNA was sequenced on platform Illumina NovaSeq 6000 (Illumina, San Diego, CA, USA). The software FastQC (version: 0.20.1) was used to filter the original reads and discard the low-quality reads, generating 88.68Gb clean data (Cock et al., 2010).

Similarly, a SMRT cell sequencing library containing about 15-20 kb fragment was constructed and sequenced using PacBio sequel II sequencing platform. In total, 81.79 Gb clean data were obtained.

For the Hi-C library, young leaves of the same *D. cambodiana* plant was selected and crosslinked by 2% formaldehyde. The chromatin was extracted from the fixed tissues and digested by DpnII endonuclease. The DNA were recovered with biotin-labeled nucleotides and the resulting blunt endswere ligated to each other using DNA ligase. DNA was purified by phenol-chloroform extraction and sheared to a size of 200-600bp fragments by sonication. Biotin labeled Hi-C sample were specifically enriched using streptavidin C1 magnetic beads. At last, the Hi-C libraries were amplified by 12-14 cycles PCR, and sequenced in Illumina HiSeq platform.

### Genome assembly and quality assessments

All subreads from SMRT sequencing, which removed sequencing connectors and low-quality sequences, were used for *D. cambodiana* genome assembly. The draft genome assembly was obtained using hifiasm software (version: 0.14.2) with the default parameters. After the sequencing data of Hi-C was filtered, a total of 196.32Gb clean data was obtained. In order to construct the chromosome level genome, we employed ALLHIC (version : 0.9.8) software with agglomerative hierarchical clustering to cluster, reorder and orient the contigs into pseudochromosomes using filtered Hi-C reads (Zhang et al., 2019). The software HiCExplorer (version: 3.6) was used for plotting based on the relationship between contig interaction intensity and position (Wolff et al. 2020). In total, 1.06 Gb of sequence was anchored into 20 pseudomolecules. Above sequencing was performed in the Wuhan Benagen Technology Co., Ltd (Wuhan, China).

### Genome annotation

Firstly, software RepeatModeler (version: open-1.0.11) was used to predict the model sequence based on its genome sequence based on denovo method, and LTR_FINDER (version: Official release of LTR_FINDER_parallel) was used to predict the LTR sequence (Smit et al., 2015; Ellinghaus et al., 2008). The software LTR_retriever (version: v2.9.0) was used to deredundant the result sequence predicted by LTR_FINDER to obtain the non-redundant LTR sequence. The two denovo sequences were merged to obtain the denovo repeat sequence library. After combining RepBase (version: 20181026) library and denovo library, software RepeatMasker (version: open-4.0.9) was used to compare and predict the repeat sequence to get the De novo + repbase result (Tarailo-Graovac et al., 2009). The RepeatMasker software RepeatProteinMask was used to predict the TE_proteion type repeat sequence, and the TE protiens result was obtained. Finally, all the repeated prediction results were Combined to get the Combined TEs result of the final genome repeat set.

For second-generation transcriptome prediction, the original reads were filtered using fastp software (version: 0.21.0) (Chen et al., 2018). The filtered reads were compared using hisat2 software (version: 2.1.0) (Kim et al., 2019), and the resulting transcript of the bam file was reconstructed through stringtie (version: 2.1.4) (Kovaka et al., 2019). Then, TransDecoder (version: v5.1.0) is used to predict the coding frame of the predicted transcript region, and the predicted coding gene is finally obtained (Haas et al., 2013). In the third-generation sequencing, ccs sequences were identified by PacBio full-length analysis using ccs in smrtlink (version: 11.0), and then isoseq3 in smrtlink was used for full-length identification, internal error correction and clustering, and the software cd-hit was further used for redundancy analysis. The full-length redundancy identified was removed, and the full-length sequence after redundancy was corrected and compared with the genome using minimap2 software (version:2.17-r941) (Li 2016). The result was compared with the transcript reconstructed by stringtie (version: 2.1.4) of bam file, and then the predicted transcript region was predicted by TransDecoder software (version: v5.1.0) for coding frame (Haas et al., 2013; Kovaka et al. 2019). Finally, the predicted coding gene is obtained. The subsequent analysis was carried out after combining the bam obtained from the two analyses, and the analysis method was the same as that of the second generation.

The protein sequences encoded by genes in the gene set were compared with the existing protein databases Uniprot (version: 202011), NR (version:202011) and the metabolic pathway database KEGG by diamond blastp to obtain the functional information of the sequences and the metabolic pathway information that the proteins may participate in (Apweiler et al. 2004; Deng et al. 2006; Kanehisa and Goto 2000). Searches for gene motifs and domains were performed using InterProScan (version: 5.52-86.0), Pfam (version: 202011) and HMMER (version:3.3.2) (Blum et al. 2021; Finn et al. 2014). The GO terms (version:202011) for genes were used. Pathway annotation was performed using KOBAS (version: v3.0) against the KEGG database (Xie et al. 2011).

For non-coding RNA annotation, tRNAscan-SE (version: 1.23) was used to search for tRNA sequences in the genome (Chan et al. 2021). rRNA prediction using rRNA database; INFERNAL (version: 1.1.2) was used to searche ncRNA sequences in the genome (Nawrocki and Eddy 2013).

### Gene family construction and phylogenetic analysis

Based on all amino acid sequences of the selected species, orthofinder software was used (version: 2.3.12) for gene family clustering, where blastp (version: 2.6.0) for comparison (Camacho et al. 2009; Emms and Kelly 2019).

Muscle (version: 3.8.31) was used to perform multiple sequence alignment of protein sequences for each single-copy gene family, and trimal (version: 1.2rev59) filter the comparison results, and then merge the filtered comparison results and connect them to super gene (Capella-Gutiérrez et al., 2009; Robert and Edgar 2004). Finally, RAxML software (version: 8.2.10) was used to construct ML species phylogenetic tree based on supergene (Alexandros n.d.). The number of strict single copy genes in this analysis was 367.

Species divergence times were estimated by PAML (version: v.4.9) and calibrated from TimeTree database(http://www.timetree.org/) (Yang 2007).

Expanded and contracted gene families were determined by Computational Analysis of gene Family Evolution (CAFÉv.3.1) (Han et al. n.d.).

### WGD

By comparing protein sequences of different species using BLAST (version: 2.6.0+), and then analyzing genome co-linearity blocks using MCScanX (version:0.8;website: https://github.com/wyp1125/MCScanx), the synonymous mutation frequency (Ks), non-synonymous mutation frequency (Ka), and the ratio of non-synonymous to synonymous mutations (Ka/Ks) for co-linear gene pairs were calculated using the yn00 module in PAML (version: 4.9) (Camacho et al., 2009; Wang et al., 2012; Yang 2007). Plot density using ggplot2 (version: 2.2.1).

## FOUNDS

This work was supported by the program of the CAMS Initiative for Innovative Medicine (2021-1-I2M-032), the National Natural Science Foundation of China (82173925), the National Science Foundation of Beijing (7222286),and the Natural Science Foundation of Hainan Province of China(322QN337).

## Author Contributions

XY and PJ conceived and designed the project, and participated in all the related analyses. XY and PJ performed the data analyses and drafted the manuscript. PJ, LS and LY contributed to the sample preparation used for genome and RNA sequencing.All authors contributed to the writing of the paper.

## Data Availability Statement

The raw data of genome and transcriptome sequencing reported in this paper will be deposited in the Genome Sequence Archive in BIG Data Center, Beijing Institute of Genomics (BIG), Chinese Academy of Sciences under the accession number (PRJCA027373). It will be publicly accessible at (http://bigd.big.ac.cn/gsa) when paper is accepted.

## ACKNOWLEDGMENTS

The authors declare no competing financial interests.

## Supplementary

**Figure S1.**
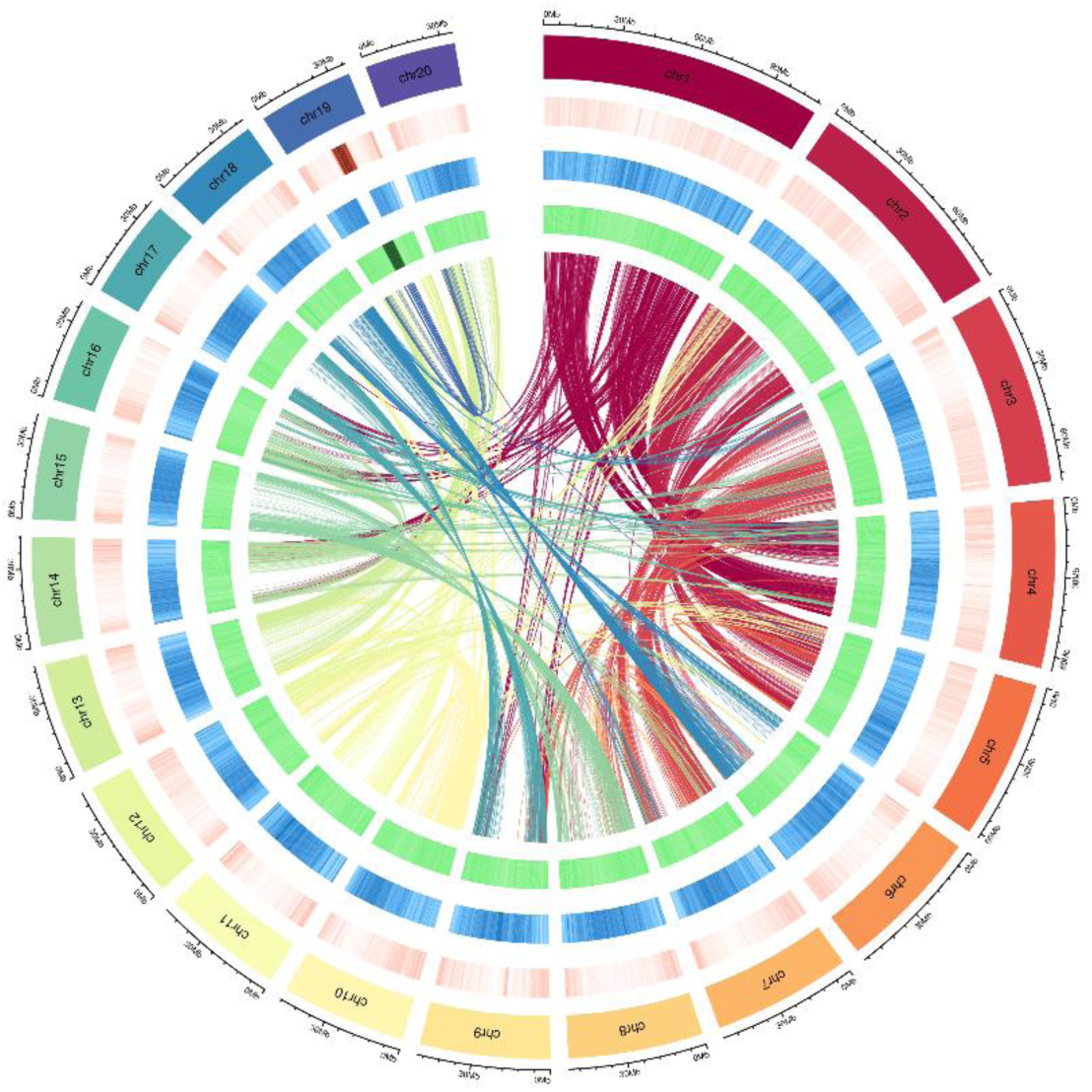
Genomic features of *D. cambodiana*. From the outer circle to the inner circle, gene density, repeat density, and GC content are calculated according to the window of 50kbp, respectively.

**Figure S2.**
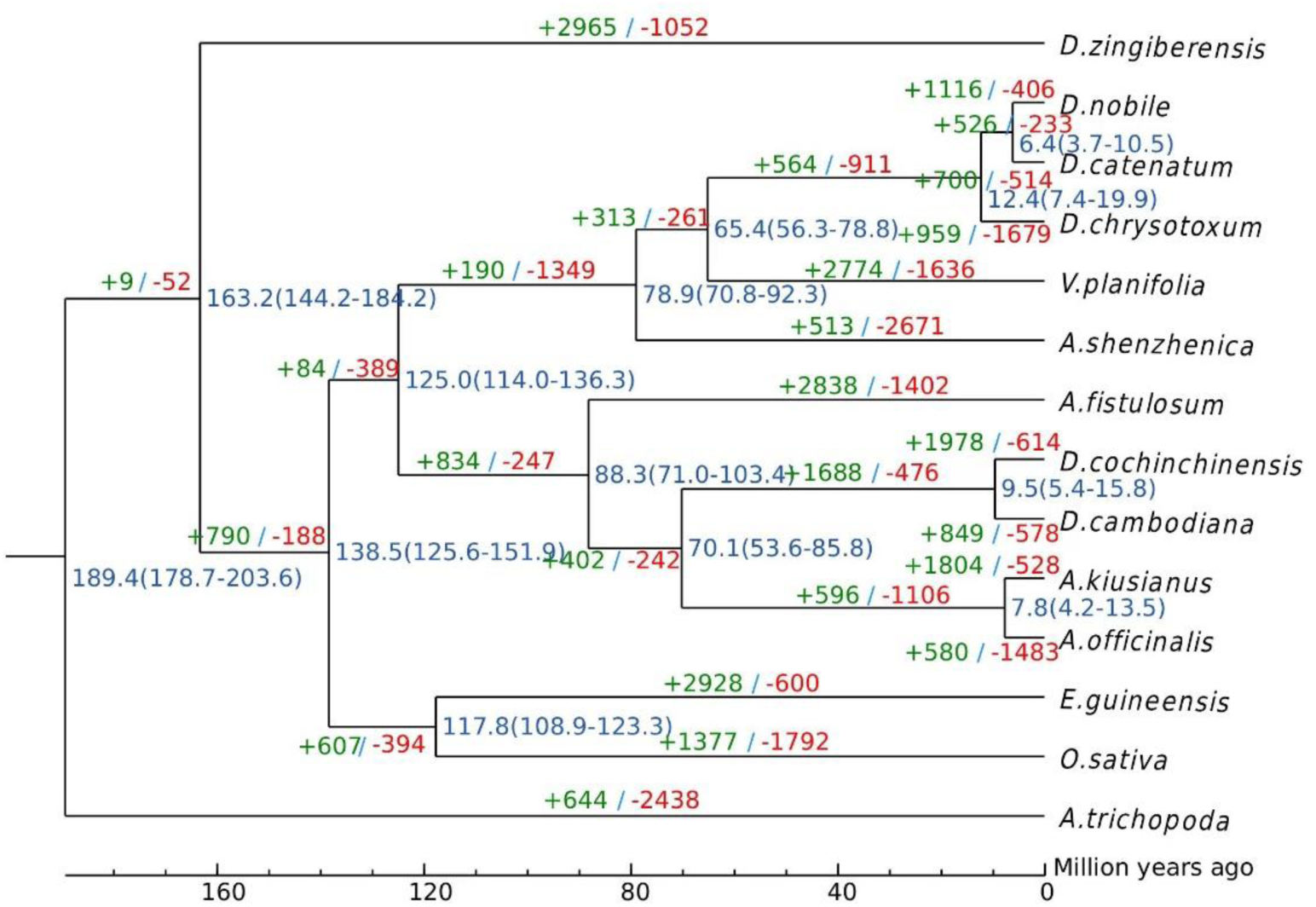
Phylogenetic tree of 14 plant species. Divergence time (Mya) estimates are indicated by the blue numbers beside the branch nodes. The numbers of gene-family contractions and expansions are indicated by green and red numbers, respectively

**Figure S3.**
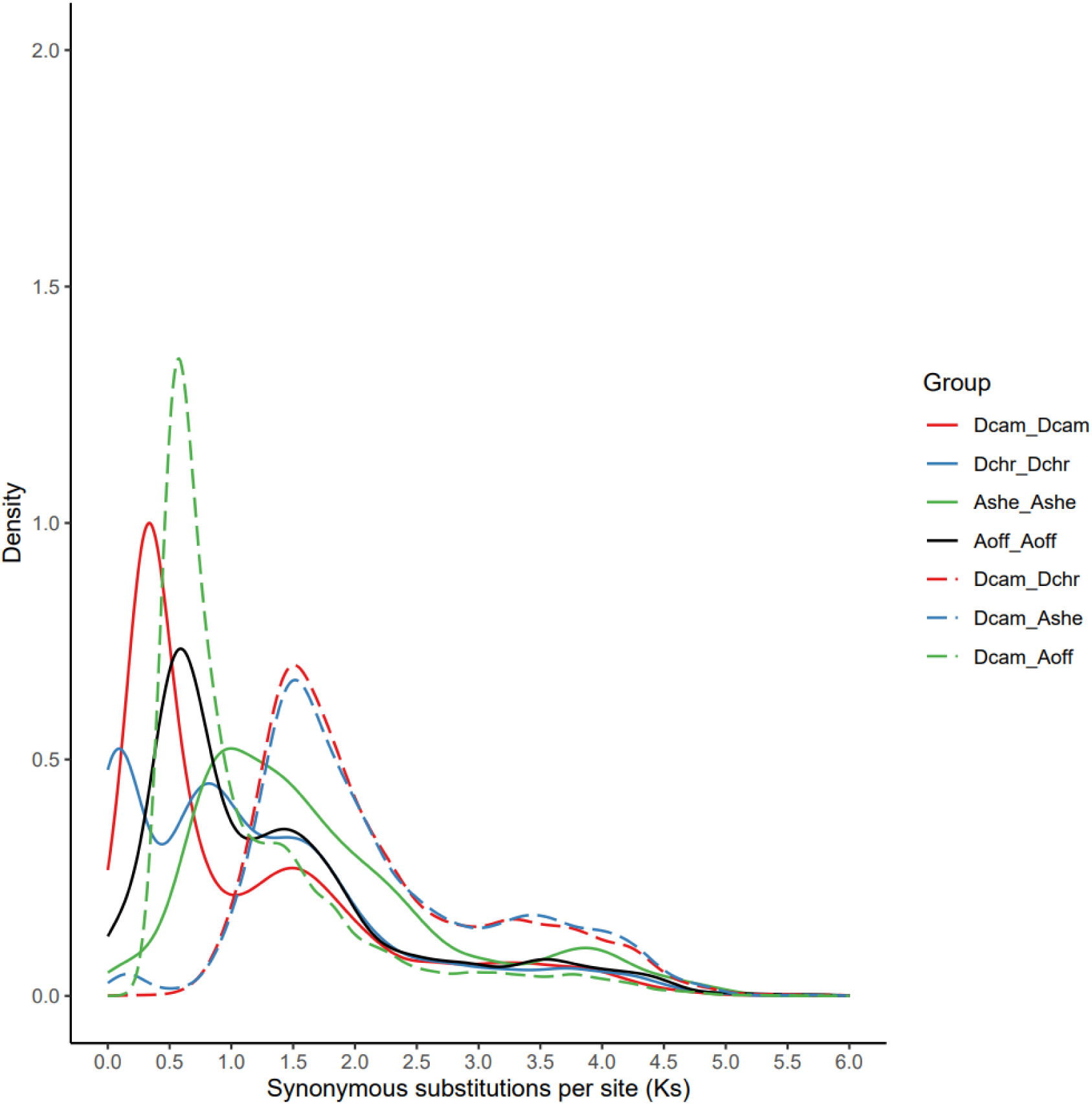
Distribution of Ks for paralogous genes among *D. cambodiana* and three other representative plant species (*D. cochinchinensis*, *A. officinalis*, and *A. shenzhenica*).

**Figure S4.**
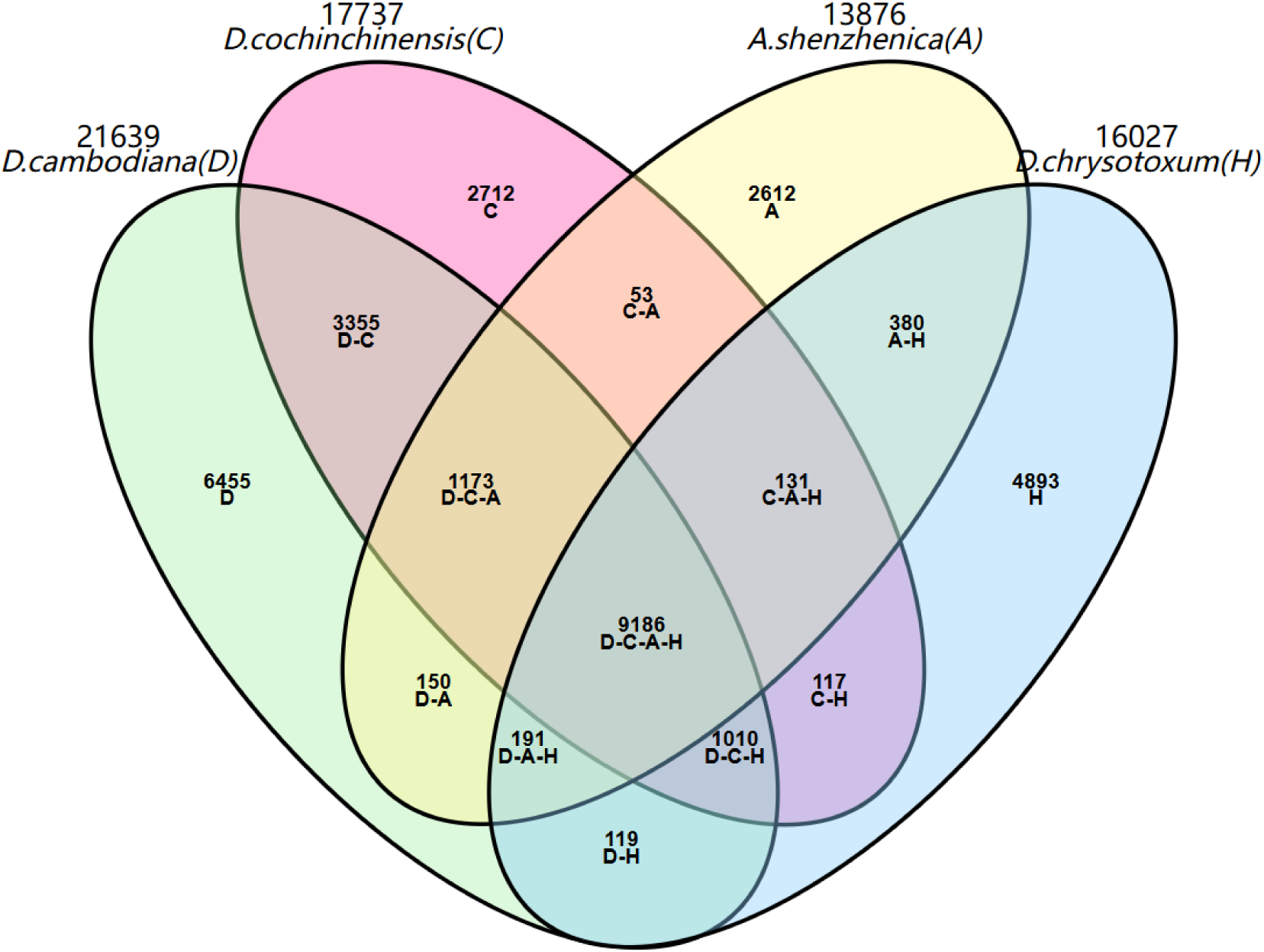
Statistical Venn diagram of gene family clustering results of four species Numbers indicate the number of gene families; The letters in parentheses are abbreviations of the Latin name of the species; The abbreviated linked strings represent the gene families shared by these species; A single acronym indicates the number of gene families unique to each of the four species.

**Figure S1.**
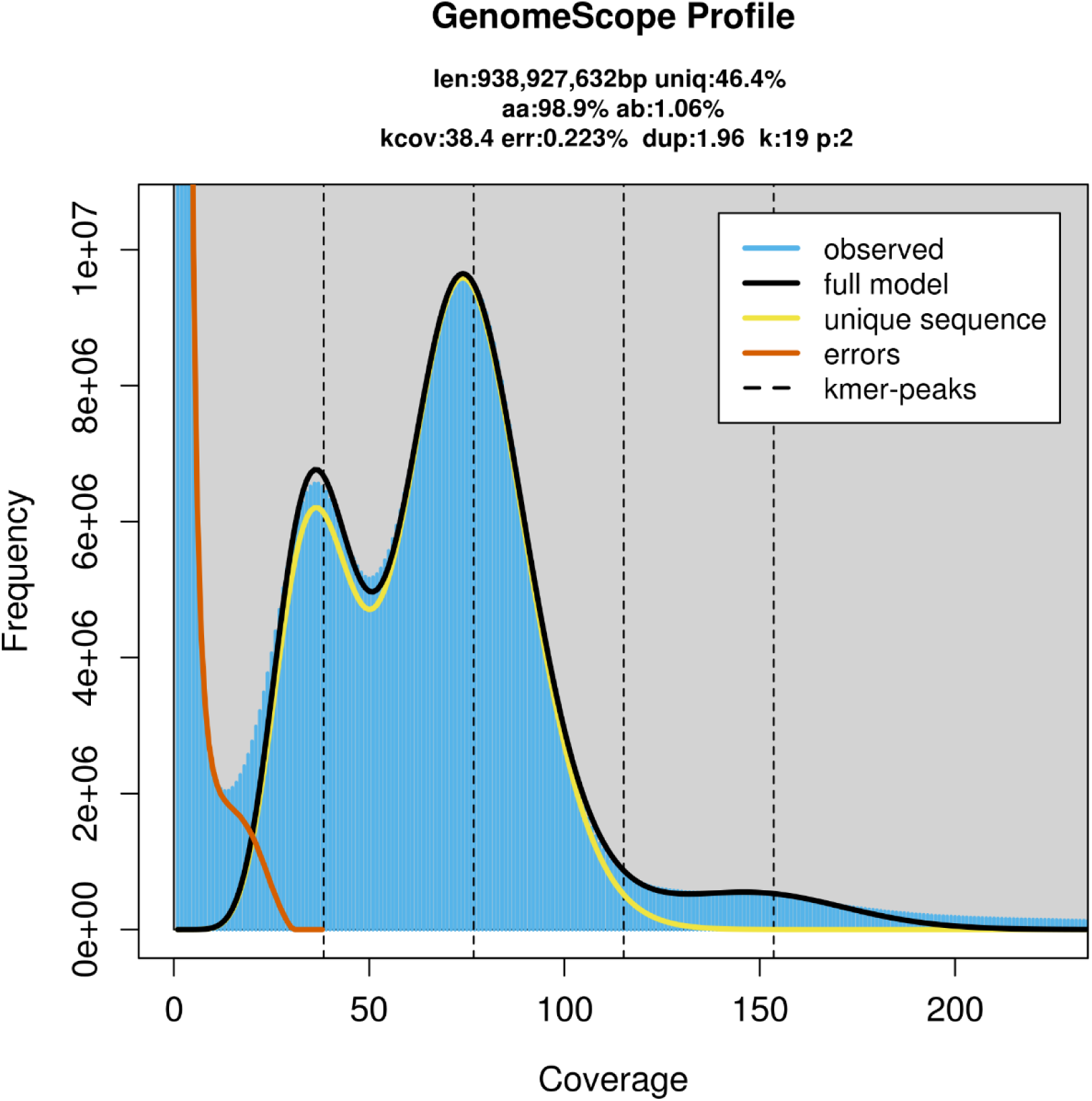
Evaluation of the genome size of *Dracaena cambodiana*. The distribution of 19-mer depth of high-quality reads was used to estimate the genome size of *D. cambodiana*. Approximately, 88.68 Gb of sequencing reads from short-insert libraries were selected and then split into 19-bp sequences (19-mers) to plot the frequency (depth) of the 19-mers.

**Figure S2.**
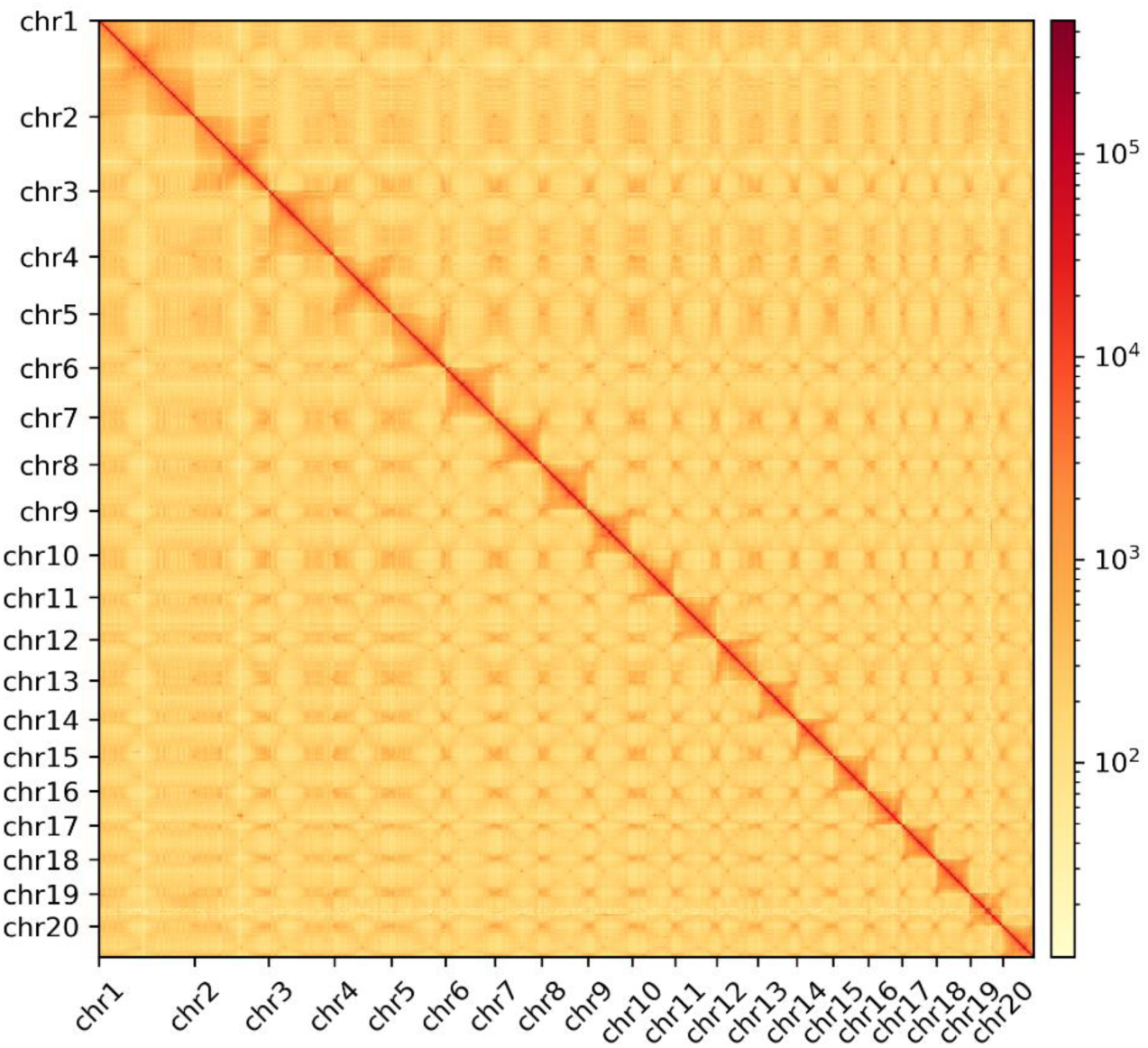
Diagram of interzone interactions within chromosomes of *Dracaena cambodiana*.

**Figure S3A.**
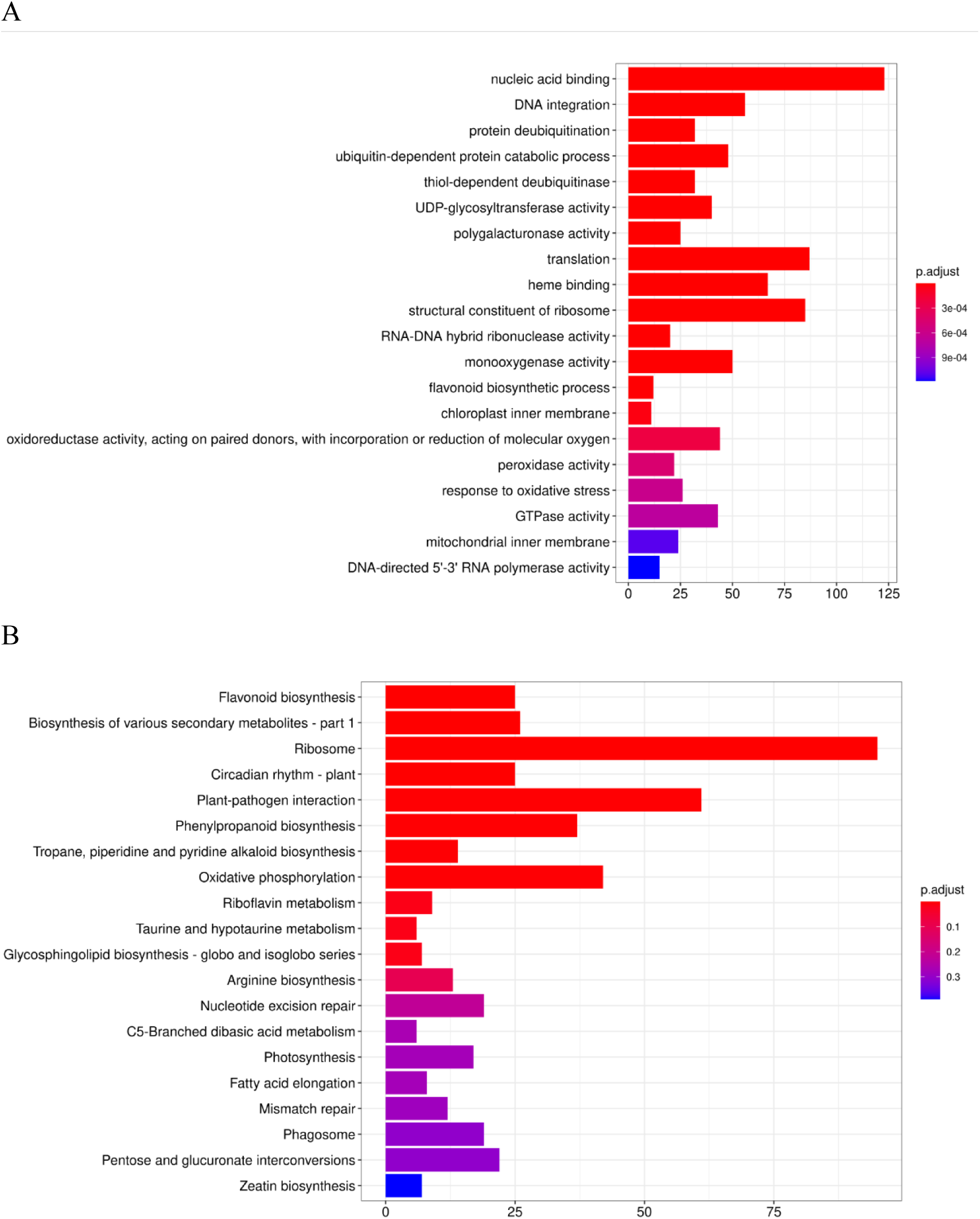
GO enrichment results of extended gene family S3B. KEGG enrichment results of extended gene family

**Figure S4A.**
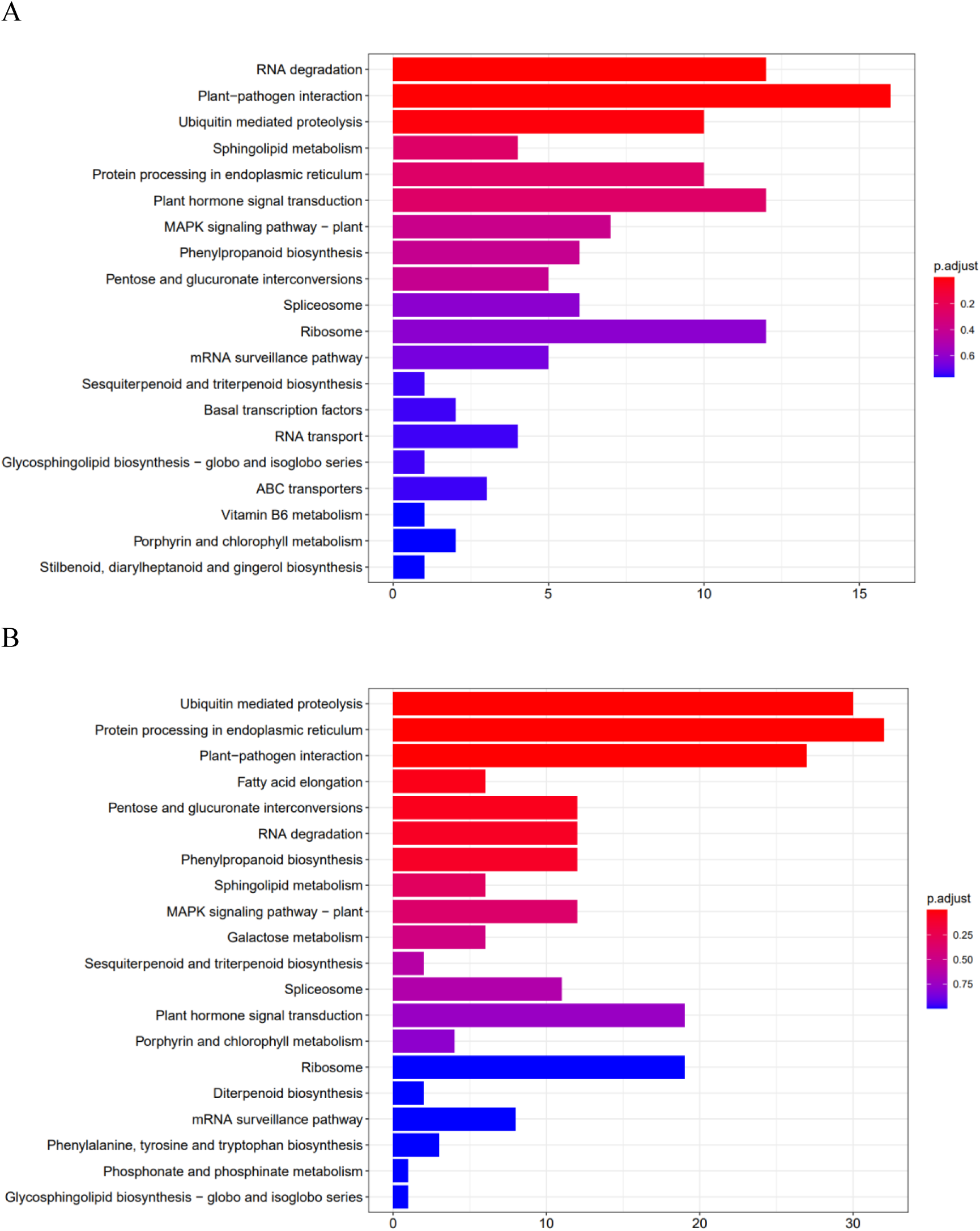
KEGG enrichment results of common genes of *D. cambodiana*. S4B. KEGG enrichment result of common genes of *D. cochinchinensis*

## References

Alexandros, S. (n.d.). “RAxML version 8: A tool for phylogenetic analysis and post-analysis of large phylogenies,” Bioinformatics, 1312–1313.

Apweiler, R., Bairoch, A., Wu, C. H., Barker, W. C., Boeckmann, B., Ferro, S., Gasteiger, E., Huang, H., Lopez, R., Magrane, M., and others (2004), “UniProt: The universal protein knowledgebase,” Nucleic acids research, Oxford University Press, 32, D115–D119.

Ashburner, M., Ball, C. A., Blake, J. A., Botstein, D., Butler, H., Cherry, J. M., … & Sherlock, G. (2000). Gene ontology: tool for the unification of biology. Nature genetics, 25(1), 25–29.

Blum, M., Chang, H.-Y., Chuguransky, S., Grego, T., Kandasaamy, S., Mitchell, A., Nuka, G., Paysan-Lafosse, T., Qureshi, M., Raj, S., and others (2021), “The InterPro protein families and domains database: 20 years on,” Nucleic acids research, Oxford University Press, 49, D344–D354.

Bos, J. Dracaena. (1998) In the Families and Genera of *Vascular Plants*; Kubitzki, K., Ed.; Springer: Berlin/Heidelberg, Germany, Volume III, pp. 238–241.

Camacho, C., Coulouris, G., Avagyan, V., Ma, N., Papadopoulos, J., Bealer, K., and Madden, T. L. (2009), “BLAST+: Architecture and applications,” BMC bioinformatics, Springer, 10, 1–9.

Capella-Gutiérrez, S., Silla-Martínez, J. M., and Gabaldón, T. (2009), “trimAl: A tool for automated alignment trimming in large-scale phylogenetic analyses,” Bioinformatics, Oxford University Press, 25, 1972–1973.

Chan, P. P., Lin, B. Y., Mak, A. J., and Lowe, T. M. (2021), “tRNAscan-SE 2.0: Improved detection and functional classification of transfer RNA genes,” BioRxiv, Cold Spring Harbor Laboratory, 614032.

Chen, S., Zhou, Y., Chen, Y., and Gu, J. (2018), “Fastp: An ultra-fast all-in-one FASTQ preprocessor,” Bioinformatics, Oxford University Press, 34, i884–i890.

Cock, P. J., Fields, C. J., Goto, N., Heuer, M. L., & Rice, P. M. (2010). The Sanger FASTQ file format for sequences with quality scores, and the Solexa/Illumina FASTQ variants. Nucleic acids research, 38(6), 1767–1771.

De Bie, T., Cristianini, N., Demuth, J.P., and Hahn, M.W. (2006). CAFE: a computational tool for the study of gene family evolution.Bioinformatics 22:1269–1271.

Deng, Y., Li, J., Wu, S., Zhu, Y., Chen, Y., He, F., and others (2006), “Integrated nr database in protein annotation system and its localization,” Comput Eng, 32, 71– 2.

Ding, X., Zhu, J., Wang, H., Chen, H., and Mei, W. (2020). Dragon’s blood from *Dracaena cambodiana* in China: applied history and induction techniques toward formation mechanism. Forests 11:372.

Edwards, H.G.M., Farwell, D.W., and Quye, A. (1997). ‘Dragon’s blood’ characterization of an ancient resin using Fourier transform Raman spectroscopy. J. Raman Spectrosc. 28:243–249.

Ellinghaus, D., Kurtz, S., & Willhoeft, U. (2008). LTRharvest, an efficient and flexible software for de novo detection of LTR retrotransposons. BMC bioinformatics, 9, 1–14.

Emms, D. M., and Kelly, S. (2019), “OrthoFinder: Phylogenetic orthology inference for comparative genomics,” Genome biology, 20.

Finn, R. D., Bateman, A., Clements, J., Coggill, P., Eberhardt, R. Y., Eddy, S. R., Heger, A., Hetherington, K., Holm, L., Mistry, J., and others (2014), “Pfam: The protein families database,” Nucleic acids research, Oxford University Press, 42, D222–D230.

Govaerts, R.; Zonneveld, B.J.M.; Zona, S.A. (2019) World checklist of Asparagaceae. Facilitated by the Royal Botanic Gardens, Kew. Available online: http://apps.kew.org/wcsp/ (accessed on 22 December 2019).

Haas, B. J., Papanicolaou, A., Yassour, M., Grabherr, M., Blood, P. D., Bowden, J., … & Regev, A. (2013). De novo transcript sequence reconstruction from RNA-seq using the Trinity platform for reference generation and analysis. Nature protocols, 8(8), 1494–1512.

Han, M. V., Thomas, G. W., Lugo-Martinez, J., & Hahn, M. W. (2013). Estimating gene gain and loss rates in the presence of error in genome assembly and annotation using CAFE 3. Molecular biology and evolution, 30(8), 1987–1997.

Hao, Q., Saito, Y., Matsuo, Y., Li, H.Z., and Tanaka, T. (2015). Chalcanestilbene conjugates and oligomeric flavonoids from Chinese dragon’s blood produced from Dracaena cochinchinensis. Phytochemistry 119:76–82.

Hubálková, I., Maděra, P., Volařík, D. (2015) Growth dynamics of *Dracaena cinnabari* under controlled conditions as the most effective way to protect endangered species. Saudi J Biol. Sci. 24(7), 1445–1452.

IUCN red list of threatened species version 2017.2. Available online: http://www.iucnredlist.org (accessed on 25 October 2017).

Kamel, M., Ghazaly, U.M., Callmander, M.W. (2018). Conservation status of the endangered Nubian dragon tree *Dracaena ombet* in Gebel Elba national park. Egypt. Oryx 49, 704–709.

Kanehisa, M., Furumichi, M., Sato, Y., Kawashima, M., & Ishiguro-Watanabe, M. (2023). KEGG for taxonomy-based analysis of pathways and genomes. Nucleic acids research, 51(D1), D587–D592.

Kanehisa, M., and Goto, S. (2000), “KEGG: Kyoto encyclopedia of genes and genomes,” Nucleic acids research, Oxford University Press, 28, 27–30.

Kim, D., Paggi, J. M., Park, C., Bennett, C., and Salzberg, S. L. (2019), “Graph-based genome alignment and genotyping with HISAT2 and HISAT-genotype,” Nature biotechnology, Nature Publishing Group, 37, 907–915.

Kovaka, S., Zimin, A. V., Pertea, G. M., Razaghi, R., Salzberg, S. L., and Pertea, M. (2019), “Transcriptome assembly from long-read RNA-seq alignments with StringTie2,” Genome biology, BioMed Central, 20, 1–13.

Lang G.Z., Li C.J., Gaohu T.Y., Li C., Ma J., Yang J.Z., Zhou T.T., Yuan Y.H., Ye F., Wei J.H., Zhang D.M. (2020) Bioactive flavonoid dimers from Chinese dragon’s blood, the red resin of *Dracaena cochinchinensis*. Bioorg Chem, 97: 103659.

Li, H. (2016), “Minimap and miniasm: Fast mapping and de novo assembly for noisy long sequences,” Bioinformatics, Oxford University Press, 32, 2103–2110.

Liu, Y., Zhao, X., Yao, R., Li, C., Zhang, Z., Xu, Y., & Wei, J. H. (2021). Dragon’s blood from dracaena worldwide: Species, traditional uses, phytochemistry and pharmacology. The American Journal of Chinese Medicine, 49(06), 1315–1367.

Luo, Y., Wang, H., Zhao, Y.X., Zeng, Y.B., Shen, H.Y., Dai, H.F., and Mei, W.L. (2011). Cytotoxic and antibacterial flavonoids from dragon’s blood of Dracaena cambodiana. Planta Med. 77:2053–2056.

Mabberley, D.J. (2008) Mabberley’s Plant-Book, 3rd ed.; Cambridge University Press: New York, NY, USA, p. 1021.

Nawrocki, E. P., and Eddy, S. R. (2013), “Infernal 1.1: 100-fold faster RNA homology searches,” Bioinformatics, Oxford University Press, 29, 2933–2935.

Pang, D.R., Su, X.Q., Zhu, Z.X., Sun, J., Li, Y.T., Song, Y.L., Zhao, Y.F., Tu, P.F., Zheng, J., and Li, J. (2016). Flavonoid dimers from the total phenolic extract of Chinese dragon’s blood, the red resin of Dracaena cochinchinensis. Fitoterapia 115:135–141.

Pruitt, K. D., Tatusova, T., & Maglott, D. R. (2005). NCBI Reference Sequence (RefSeq): a curated non-redundant sequence database of genomes, transcripts and proteins. Nucleic acids research, 33(suppl_1), D501–D504.

Robert, C., and Edgar (2004), “MUSCLE: Multiple sequence alignment with high accuracy and high throughput.” Nucleic acids research.

Simaõ, F.A., Waterhouse, R.M., Ioannidis, P., Kriventseva, E.V., and Zdobnov, E.M. (2015). BUSCO: assessing genome assembly and annotation completeness with single-copy orthologs. Bioinformatics 31:3210–3212.

Smit, A., Hubley, R., and Green, P. (2015), “RepeatModeler open-1.0 (2008–2015),” Seattle, USA: Institute for Systems Biology. Available from: http://www.repeatmasker.org, Last Accessed May, 1, 2018.

Spampinato, C. P. (2017). Protecting DNA from errors and damage: an overview of DNA repair mechanisms in plants compared to mammals. Cellular and Molecular Life Sciences, 74, 1693–1709.

Sun HF, Song MF, Zhang Y, Zhang ZL. (2021) Transcriptome profiling reveals candidate flavonoid-related genes during formation of dragon’s blood from *Dracaena cochinchinensis* (Lour.) S.C.Chen under conditions of wounding stress. Journal of Ethnopharmacology, 273, 113987.

Tarailo-Graovac, M., & Chen, N. (2009). Using RepeatMasker to identify repetitive elements in genomic sequences. Current protocols in bioinformatics, 4-10.

Tatusov, R. L., Fedorova, N. D., Jackson, J. D., Jacobs, A. R., Kiryutin, B., Koonin, E. V., … & Natale, D. A. (2003). The COG database: an updated version includes eukaryotes. BMC bioinformatics, 4, 1–14.

Tomlinson, P.B., and Huggett, B.A. (2012). Cell longevity and sustained primary growth in palm stems. Am. J. Bot. 99:1891–1902.

Wang, Y., Tang, H., Debarry, J. D., Tan, X., Li, J., Wang, X., Tae-Ho, L., Jin, H., Barry, M., and Guo, H. (2012), “MCScanX: A toolkit for detection and evolutionary analysis of gene synteny and collinearity,” Nucleic Acids Research, 40, e49–e49.

Wolff, J., Rabbani, L., Gilsbach, R., Richard, G., Manke, T., Backofen, R., and Grüning, B. A. (2020), “Galaxy HiCExplorer 3: A web server for reproducible hi-c, capture hi-c and single-cell hi-c data analysis, quality control and visualization,” Nucleic acids research, Oxford University Press, 48, W177–W184.

Xie, C., Mao, X., Huang, J., Ding, Y., Wu, J., Dong, S., Kong, L., Gao, G., Li, C.-Y., and Wei, L. (2011), “KOBAS 2.0: A web server for annotation and identification of enriched pathways and diseases,” Nucleic acids research, Oxford University Press, 39, W316–W322.

Xin Yaxuan, Tan Yunhong, Xin Peiyao, Li Haitao, Zhang Shuhong, Tang Junrong, Yu Wenbin. (2023) Taxonomical clarification of *Dracaena cambodiana* Pierre ex Gagnep., the source plant of Chinese “Resina Draconis”. Plant Science Journal, DOI: 10.11913/PSJ. 2095-0837. 23340.

Xu Y., Zhang K., Zhang Z., Liu Y., Lv F., Sun P., Gao S., Wang Q., Yu C., Jiang J., Li C., Song M., Gao Z., Sui C., Li H., Jin Y., Guo X., and Wei J. (2022). A chromosome-level genome assembly for *Dracaena cochinchinensis* reveals the molecular basis of its longevity and formation of dragon’s blood. Plant Comm. 3, 100456.

Yang L.R., Zhang Z.L., Yun Y., Yan W.P., Chen X., Zhang L., Zheng D.J., Chen J. L. (2018) The population structure and dynamics of *Dracaena cambodiana*, an endangered tree on Hainan Island. Acta Ecologica Sinica, 38(8): 2802–2815

Yang, Z. (2007), “PAML 4: Phylogenetic analysis by maximum likelihood,” Molecular Biology and Evolution, 24, 1586–1591.

Zhang, X., Zhang, S., Zhao, Q., Ming, R., and Tang, H. (2019), “Assembly of allele-aware, chromosomal-scale autopolyploid genomes based on hi-c data,” Nature plants, Nature Publishing Group, 5, 833–845.

Zheng, D.J., Xie, L.S., Wang, Y., Zhang, Z.L., and Zhang, W. (2009). Research advances in dragon’s blood plants in China. Chin. Wild Plant Res. 28:15–20.

Zhu JH, CaoTJ, Dai HF, Li HL, Guo D, Mei WL, Peng SQ. (2016) *De Novo* transcriptome characterization of *Dracaena cambodiana* and analysis of genes involved in flavonoid accumulation during formation of dragon’s blood. Scientific Reports, 6: 38315.

